# Lipid Scrambling Amplifies Plasma Membrane Undulations via Anti-Registered Domains Stabilized by Long-Chain Sphingomyelin

**DOI:** 10.64898/2026.11.11.765848

**Authors:** Teppei Yamada, Wataru Shinoda

**Affiliations:** Graduate School of Environmental, Life, Natural Science and Technology, Okayama University, 3-1-1 Tsushima-naka, Kita-ku, Okayama 700-8530, Japan; Research Institute for Interdisciplinary Science, Okayama University, 3-1-1 Tsushima-naka, Kita-ku, Okayama 700-8530, Japan

## Abstract

Lipid scrambling, which dissipates the transbilayer lipid asymmetry of plasma membranes (PMs), plays an essential role in various large-scale membrane remodeling processes. Although lipid domains have been implicated in these processes, the underlying mechanisms remain poorly understood, owing to the lack of a molecular-level picture of how lipid scrambling reorganizes these domains. Here, using coarse-grained molecular dynamics simulations of asymmetric and fully scrambled PM models, we show that lipid scrambling markedly enhances membrane undulations. We identify two key features of the domain organization underlying this enhancement in the scrambled PM: domain anti-registration, which couples to membrane curvature through a mismatch in preferred curvature between the two leaflets, and scrambling-induced domain enlargement, which permits this curvature to extend over large length scales. Simulations with different sphingomyelin (SM) compositions demonstrate that long-chain SMs stabilize anti-registered domains and the associated membrane curvature. Together, these results suggest a physical mechanism by which lipid scrambling primes the PM for remodeling through the reorganization of lipid domains, with long-chain SMs playing a key role. We also find that the interleaflet domain organization of the scrambled PM model depends strongly on system size, shifting from anti-registration to registration as the simulation box is reduced, and we discuss the origin of this finite-size effect.

## INTRODUCTION

Plasma membranes (PMs) maintain an asymmetric distribution of lipids between their two leaflets under steady-state conditions.^1–3^ This asymmetry gives rise to leaflet-specific cholesterol partitioning,^4^ fluidity,^3,5^ and phase behavior,^6^ thereby balancing structural stability with dynamic signaling functions.^7^ Lipid scrambling, the scramblase-mediated loss of this asymmetry, is essential for various biological processes involving large-scale membrane remodeling, including blebbing,^8^ extracellular vesicle release,^9–12^ and compensatory endocytosis.^13–15^ Because these processes involve rapid morphological changes, such as the outward budding of vesicles or the inward retrieval of excess membrane area, lipid scrambling may itself facilitate membrane deformation.

Recent studies have begun to clarify the link between lipid scrambling and membrane remodeling. For example, Wang *et al*. demonstrated that the loss of asymmetry facilitates PM blebbing by reducing average lipid packing.^8^ Le *et al*. further showed that cholesterol and GPI-anchored proteins, key components of raft domains, are enriched in extracellular vesicles released upon lipid scrambling, suggesting that extracellular vesicle biogenesis is spatially coupled to lipid domains.^12^ Despite its potential importance, how lipid scrambling reorganizes membrane domains and how this reorganization facilitates membrane remodeling, remain poorly understood, primarily because nanoscale lipid domains are difficult to resolve experimentally.^16^ Indeed, although the loss of lipid asymmetry in PM-derived vesicles has been reported to stabilize ordered domains against increasing temperature,^17^ the structural organization of these domains remains elusive.

Coarse-grained (CG) molecular dynamics (MD) simulations have been widely used to elucidate PM organization, bridging the gap between the time and length scales accessible to experiments and those accessible to all-atom (AA) MD simulations.^18–21^ In particular, the SPICA force field (FF) is well suited to studying domain organization, because it accurately reproduces the distribution functions of lipid mixtures obtained from AA models, as well as cholesterol-induced phase separation in multicomponent lipid bilayers.^22–27^ The SPICA FF has also been applied to an asymmetric PM model based on the lipid composition of the red blood cell membrane, reproducing the leaflet-specific lipid packing, transbilayer cholesterol partitioning, and interleaflet stress observed in experiments and in AA MD simulations of realistic PM models.^6^

In this study, we used CG MD simulations with the SPICA FF to compare asymmetric and fully scrambled PM models, incorporating the recently proposed imbalance in phospholipid number between the two leaflets of the PM.^4^ This approach allowed us to characterize the domain reorganization induced by lipid scrambling and to elucidate its role in membrane remodeling.

## METHODS

### Asymmetric and Scrambled Plasma Membrane Models

Based on lipidomic data for human red blood cell membranes,^3,4^ we constructed an asymmetric PM model with lateral dimensions of approximately 76 nm × 76 nm, consisting of cholesterol and 11 representative phospholipid species (Table 1). The phospholipid number asymmetry between the two leaflets was taken into account, with the inner leaflet containing 1.3 times as many phospholipids as the outer leaflet. The scrambled PM was modeled as the fully scrambled limiting state, in which all lipids of the asymmetric membrane were distributed uniformly between the two leaflets; thus, the scrambled membrane was compositionally symmetric while retaining the overall lipid composition of the asymmetric membrane. Both membranes were constructed using Packmol^28^ and solvated with 0.15 M NaCl solution. In addition, we simulated smaller systems and systems in which specific lipid species were replaced with others. The detailed lipid compositions of all systems are provided in the Supporting Information.

**Table 1.** Lipid composition of the asymmetric plasma membrane model.

| Outer leaflet |  | Inner leaflet |  |
| Lipid | Number | Lipid | Number |
|---|---|---|---|
| Cholesterol | 7836 (54.7%) | Cholesterol | 2694 (24.2%) |
| 16:0 SM | 1458 (10.2%) | 16:0-18:1 PC | 648 (5.8%) |
| 24:1 SM | 1134 (7.9%) | 16:0-18:2 PC | 1620 (14.6%) |
| 24:0 SM | 972 (6.8%) | 16:0-20:4 PS | 2430 (21.9%) |
| 18:0-18:1 PC | 810 (5.7%) | 16:0-22:6 PE | 1458 (13.1%) |
| 16:0-18:2 PC | 1620 (11.3%) | 18:1-20:4 PE | 648 (5.8%) |
| 16:0-20:4 PC | 486 (3.4%) | 18:0-22:4 PEp | 1620 (14.6%) |
| Total | 14316 | Total | 11118 |

Values in parentheses indicate the mole fraction of each lipid species within the corresponding leaflet. For cholesterol, the number averaged over the final 10 μs of the simulation is shown, because cholesterol spontaneously flip-flops between the two leaflets during the simulation. Abbreviations: SM, sphingomyelin; PC, phosphatidylcholine; PS, phosphatidylserine; PE, phosphatidylethanolamine; PEp, PE plasmalogen.

### Molecular Dynamics Simulations

All simulations were performed using the SPICA FF. CG MD simulations were carried out with GROMACS version 5.1.5,^29^ modified for the SPICA FF. The parameters and CG mapping definitions of the SPICA FF are available on the official website (https://www.spica-ff.org/), and the setup tools and the modified GROMACS source code are distributed on GitHub (https://github.com/SPICA-group). The temperature and pressure were maintained at 298 K and 1 atm, respectively, using the Nosé–Hoover thermostat^30^ and the Parrinello–Rahman barostat^31^ with semi-isotropic coupling. Nonbonded Lennard-Jones (LJ) interactions were truncated at 1.5 nm, and electrostatic interactions were evaluated using the particle-mesh Ewald (PME) method^32,33^. The time step was 5 fs. The asymmetric and scrambled PM models were simulated for 40 and 45 μs, respectively.

### Analyses

#### Two-Dimensional Height Distribution of the Bilayer Midplane

Prior to analysis, each configuration was translated so that the center of all phosphate groups coincided with the origin. To calculate the two-dimensional height distribution of the bilayer midplane, the membrane plane was divided into 1 nm × 1 nm grid cells. The midplane height in each cell was defined as the average *z*-coordinate of the phosphate groups of both leaflets within that cell. These two-dimensional distributions were averaged over 10 ns intervals, and the root-mean-square (RMS) height fluctuation at each time point was evaluated from the resulting distributions.

#### Order Parameter

To characterize domain structures, we calculated the two-dimensional distribution of the orientational order parameter, *S*, of the phospholipid tails in each leaflet:

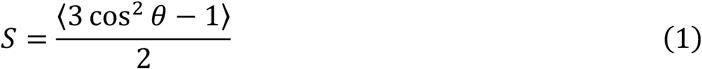

where *θ* is the angle between a CG bond vector in the lipid tail and the bilayer normal (*z*-axis), and the angular brackets denote the average over all CG bonds in both hydrocarbon chains of each phospholipid, including the sphingosine and acyl chains of SMs. This parameter equals 1 when the lipid tails are perfectly aligned with the bilayer normal and 0 when they are randomly oriented.

To compute the distribution of *S*, the average order parameter of the phospholipids was calculated in each 1 nm grid cell of each leaflet, with lipids assigned to cells according to their center-of-mass positions. The resulting distributions were averaged over 10 ns intervals, as in the height analysis. To evaluate the local arrangement of domains across the two leaflets, we then calculated the difference in the order parameter between the upper and lower leaflets, Δ*S* = *S*_upper_ − *S*_lower_. Because the two leaflets of the scrambled PM are compositionally identical, they are referred to as the upper and lower leaflets rather than the outer and inner leaflets.

#### Spatial Correlation Function of Order Parameter

To evaluate domain size, we calculated the spatial correlation function of the order parameter *S*:

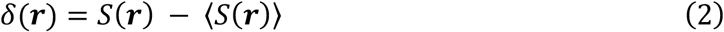

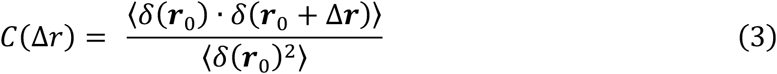

In Eq 2, ***r*** denotes a two-dimensional position vector in the bilayer plane. In Eq 3, Δ*r* on the left-hand side denotes the in-plane distance; on the right-hand side, ***r***_0_ is a reference position vector and Δ***r*** is a displacement vector of magnitude Δ*r*, with the average taken over all reference positions and displacement directions.

#### Registration Area Fraction of Lipid Domains

To determine whether lipid domains adopt a registered or anti-registered configuration, we calculated the registration area fraction. For this analysis, the grid cells of each leaflet were classified as *ordered* if their local order parameter exceeded the median value for that leaflet and as *disordered* otherwise, so that ordered and disordered cells each occupied exactly half of each leaflet. The registration area fraction was defined as the fraction of grid cells in which the two leaflets are in the same phase, that is, ordered facing ordered or disordered facing disordered. With this median-based classification, the registration area fraction is 0.5 when the domain arrangements in the two leaflets are uncorrelated, and it ranges from 0 for complete anti-registration to 1 for complete registration.

#### Mean Curvature

The local mean curvature, *H*, was calculated from the partial derivatives of the bilayer midplane height, ℎ(*x*, *y*), as follows:

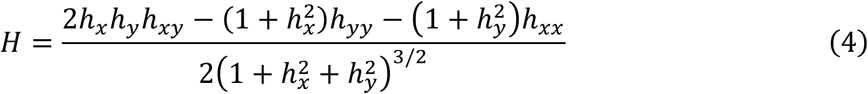

where the subscripts on ℎ denote partial derivatives of the midplane height with respect to *x* and *y*; for example, *h_x_* = ∂*h*/∂*x* and *h_xy_* = ∂^2^*h*/∂*x*∂*y*.

To suppress molecular-scale protrusion noise before differentiation, a Gaussian filter with a standard deviation of *σ* = 3 nm was applied to the midplane height distribution. As for the other properties, the curvature distributions were averaged over 10 ns intervals. The partial derivatives were evaluated numerically by central differences on the 1 nm grid. Note that the sign of *H* was inverted relative to the standard mathematical definition, following the convention for biological membranes in which an upward-convex deformation corresponds to positive curvature.

#### Correlation of Lipid Density with Δ***S*** and Mean Curvature

The two-dimensional density distribution of each lipid species was calculated in the same manner and used to evaluate the correlations of lipid density with Δ*S* and mean curvature. Because the densities of low-abundance lipids are subject to large statistical errors, a Gaussian filter with a standard deviation of *σ* = 1 nm was applied uniformly to the density distributions of all lipid species before the correlation analysis. Pearson correlation coefficients were calculated for each 10 ns-averaged snapshot over the final 10 μs of the trajectory, and their means and standard deviations were obtained from all analyzed snapshots.

## RESULTS AND DISCUSSION

### Lipid Scrambling Amplifies Plasma Membrane Undulations

The scrambled PM exhibited substantially larger undulations than the asymmetric PM, as illustrated by representative snapshots taken at the frames with the largest membrane deformation in each system (Figure 1a). To quantify these undulations, we analyzed local deviations of the bilayer midplane from its mean *z*-coordinate. Figure 1b shows the time evolution of the RMS height fluctuation of the bilayer midplane over the entire membrane plane. The RMS height fluctuation of the scrambled PM increased over time, whereas that of the asymmetric PM remained small throughout the simulation.

**Figure 1.**
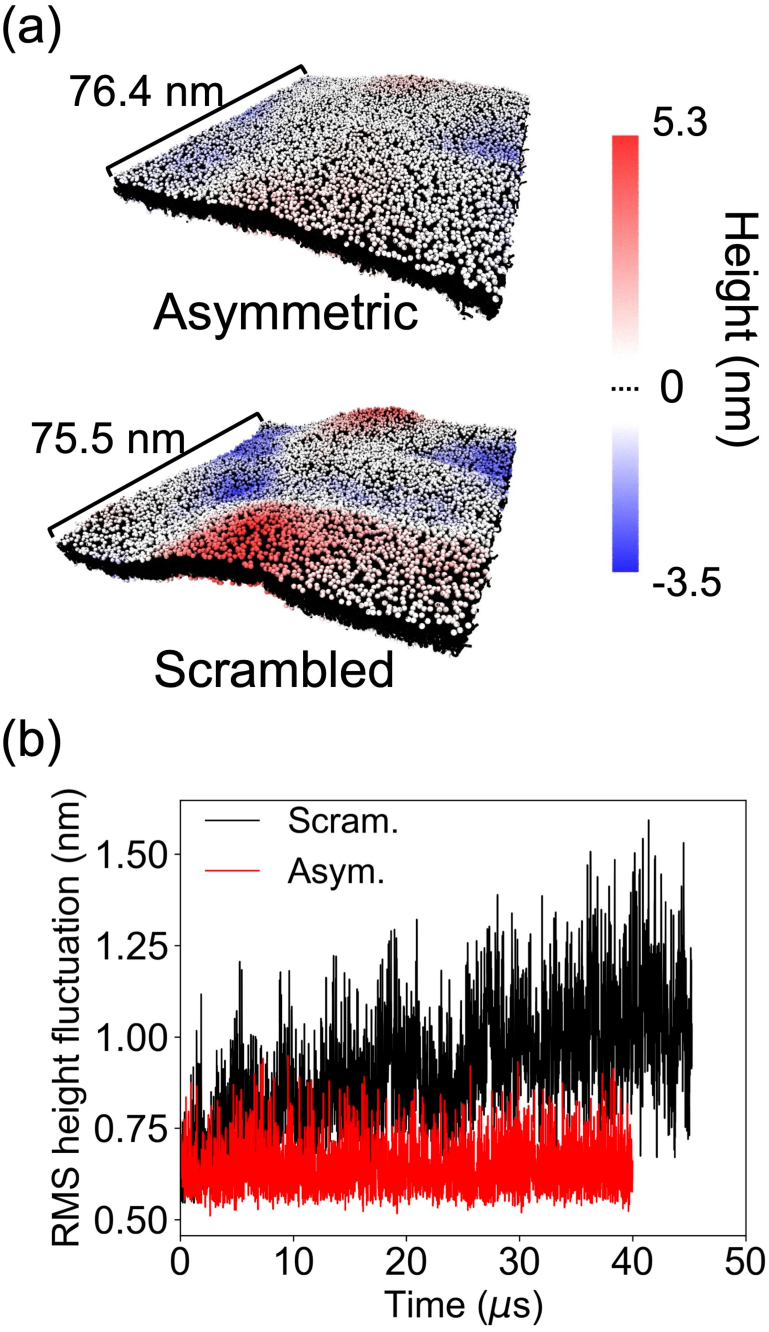
Lipid scrambling induces extensive membrane undulations. (a) Representative snapshots of the asymmetric and scrambled PMs at the frames with the largest root-mean-square height fluctuation of the bilayer midplane (see panel b). Phosphate CG beads are colored according to the deviation of their z-coordinate from the average z-coordinate of the phosphate groups in the same leaflet. (b) Time evolution of the root-mean-square height fluctuation of the bilayer midplane calculated over the entire membrane plane.

### Domain Enlargement and Anti-Registration in the Scrambled Plasma Membrane

To investigate how the enhanced undulations relate to scrambling-induced domain reorganization, we characterized domain structures through the two-dimensional distribution of the order parameter *S* in each leaflet. Figure 2 shows these distributions for both leaflets of the asymmetric (Figure 2a) and scrambled (Figure 2b) PMs, calculated from the 10 ns trajectory segments containing the frame with the largest RMS height fluctuation. In the asymmetric PM, the outer leaflet was predominantly ordered, whereas the inner leaflet was largely disordered but heterogeneous, containing small ordered domains. The functional implications of this leaflet-specific phase behavior in the asymmetric PM were discussed in detail in our recent study.^6^

In contrast, the scrambled PM exhibited distinct domains in both leaflets, with highly ordered and disordered domains (Figure 2b). The highly ordered domains were enriched in SMs and cholesterol, whereas the disordered domains were composed mainly of polyunsaturated lipids, defined here as lipids containing four or more double bonds (Figure 2c). Although lipids with two double bonds are generally classified as polyunsaturated, their preference for domains and curvature in our results was close to that of monounsaturated lipids. Thus, we grouped them together as mono-or di-unsaturated lipids.

**Figure 2.**
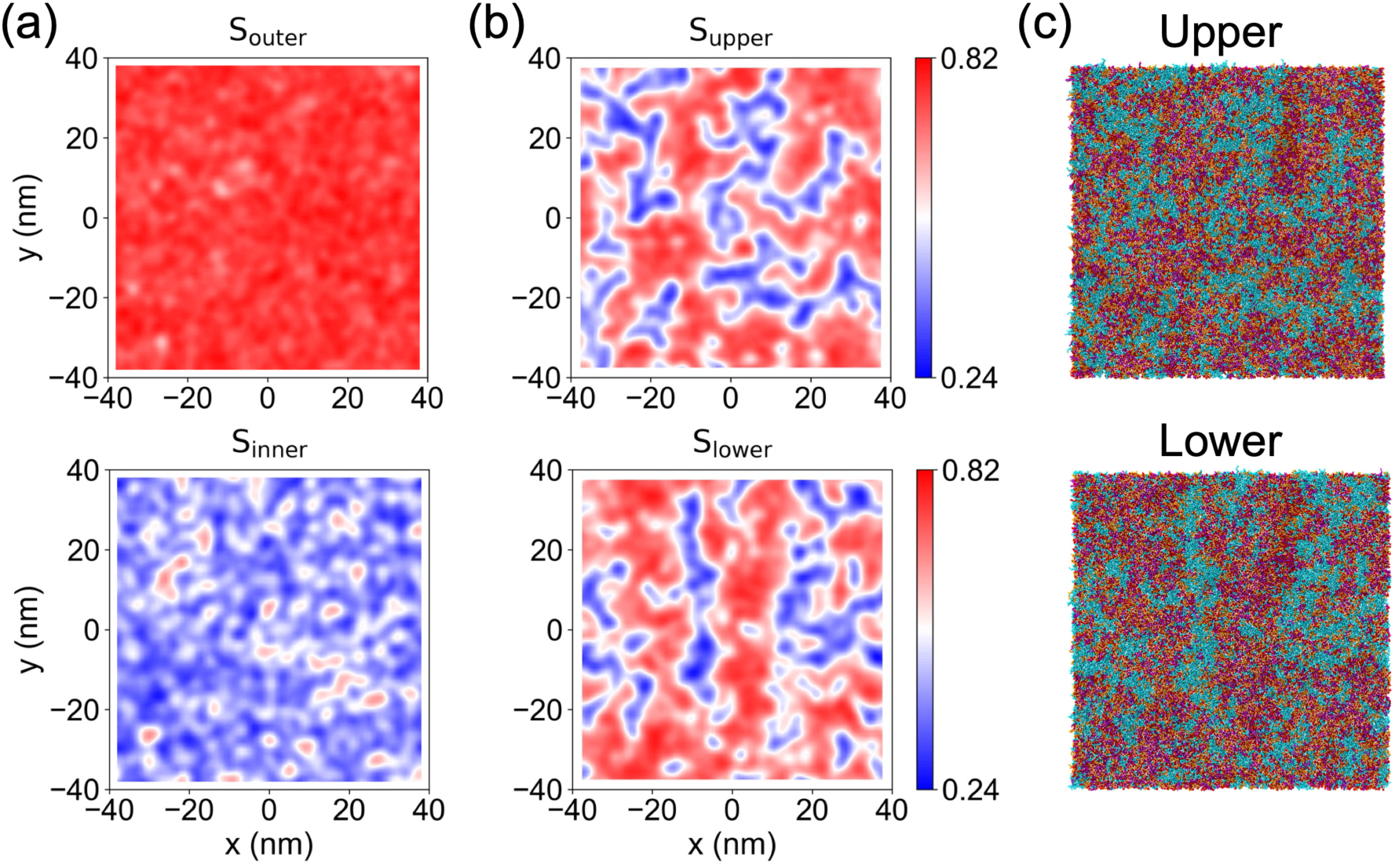
Phase behavior in the asymmetric and scrambled PMs. (a) Order parameter maps for the outer and inner leaflets of the asymmetric PM. (b) Order parameter maps for the upper and lower leaflets of the scrambled PM. (c) Top-view snapshot of the two leaflets of the scrambled PM. Color codes are as follows: sphingomyelins, purple; mono-or di-unsaturated phosphatidylcholines, orange; polyunsaturated lipids, cyan; cholesterol, red.

Notably, the domains in the scrambled PM were larger than those in the asymmetric PM. To compare the domain sizes more quantitatively, we computed the spatial correlation function of the order parameter for the outer and inner leaflets of the asymmetric PM and for the scrambled PM (averaged over the two leaflets), using the final 10 μs of each simulation (Figure 3). The correlation function of the scrambled PM decayed more slowly than those of both leaflets of the asymmetric PM, indicating a longer correlation length of lipid order. Defining the average domain radius as the distance at which *C*(Δ*r*) first crosses zero, we obtained a domain radius of 12 nm for the scrambled PM, compared with 4.0 nm for the inner leaflet of the asymmetric PM. Although this definition is not applicable to the outer leaflet of the asymmetric PM, for which *C*(Δ*r*) approaches zero only asymptotically, these results indicate that lipid scrambling enlarges domains.

**Figure 3.**
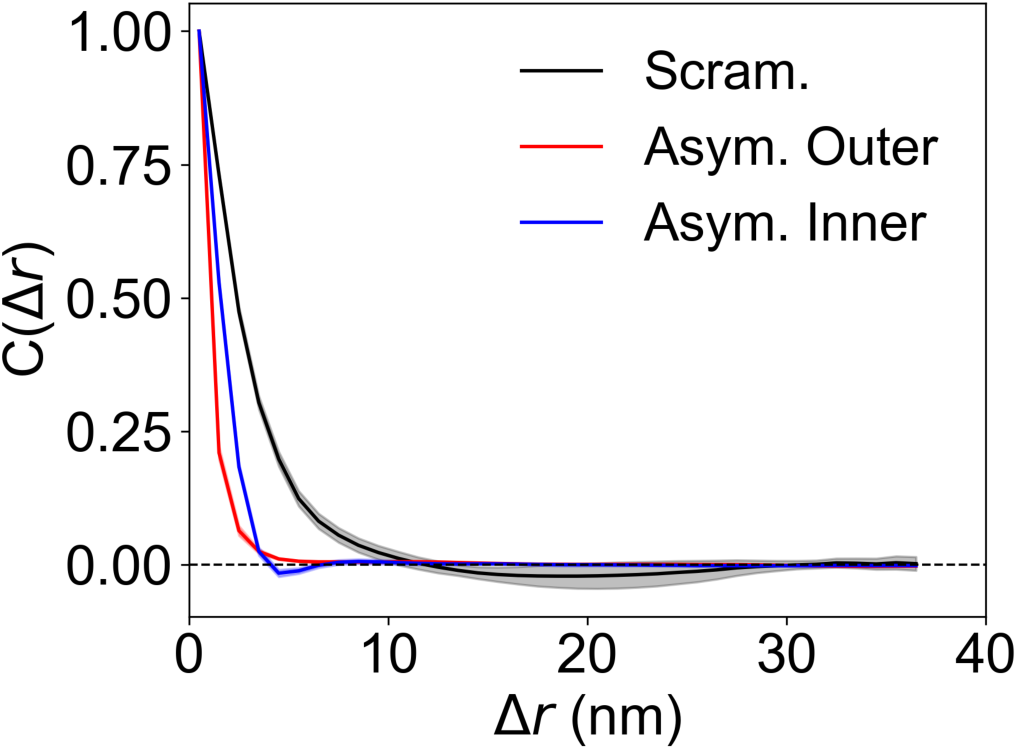
Spatial correlation function of the orientational order parameter of phospholipid tails. Correlation functions are shown for the outer and inner leaflets of the asymmetric PM and for the leaflet-averaged scrambled PM, calculated from the final 10 μs of the simulations. The shaded areas denote the standard deviations determined from five correlation functions obtained at 2 μs intervals.

Domain enlargement upon scrambling likely arises from the coexistence of high concentrations of SMs and polyunsaturated lipids within the same leaflet. In the asymmetric PM, such contacts are largely avoided because SMs are localized predominantly in the outer leaflet, which contains few polyunsaturated lipids. Upon scrambling, however, the unfavorable interaction between SMs and polyunsaturated lipids are expected to outweigh the entropy of mixing, thereby promoting the formation of larger domains.

In addition, the domains in the scrambled PM tended to adopt an anti-registered configuration, in which ordered domains in one leaflet faced disordered domains in the opposing leaflet (Figure 2b). To assess the stability of this configuration, we analyzed the time evolution of the registration area fraction (Figure 4). In the absence of an energetic preference for either the registered or the anti-registered configuration, this fraction is expected to be approximately 0.5, which was indeed the value observed in the early stage of the simulation. Thereafter, the registration area fraction remained consistently below 0.5, confirming that anti-registration is favored in the equilibrated scrambled PM. We therefore hypothesized that domain anti-registration generates a local mismatch in preferred curvature between the two leaflets, thereby inducing or amplifying membrane undulations, and that domain enlargement allows these undulations to extend over larger length scales.

**Figure 4.**
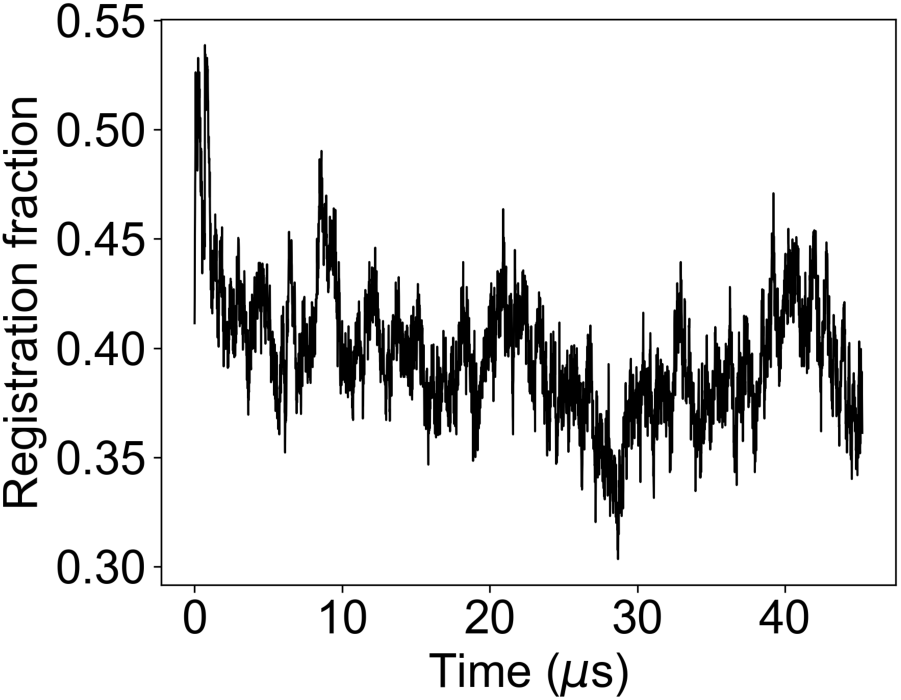
Time evolution of the registration area fraction in the scrambled PM.

### Coupling between Anti-Registered Domains and Membrane Curvature

To test this hypothesis, we calculated the two-dimensional distribution of the local mean curvature of the bilayer midplane (Figure 5a) and compared with the map of the interleaflet order parameter difference, Δ*S* = *S*_upper_ − *S*_lower_ (Figure 5b). In this map, large positive values of Δ*S* correspond to regions where highly ordered domains in the upper leaflet face disordered domains in the lower leaflet, large negative values correspond to the opposite arrangement, and values close to zero correspond to regions where domains of the same type face each other. The mean curvature and Δ*S* were strongly correlated in space: the Pearson correlation coefficient between the two distributions, calculated every 10 ns and averaged over the final 10 μs, was 0.61 ± 0.05 (Figure 5c). This correlation indicates that the coupling between domain anti-registration and local membrane curvature is a persistent feature of the equilibrated scrambled membrane.

**Figure 5.**
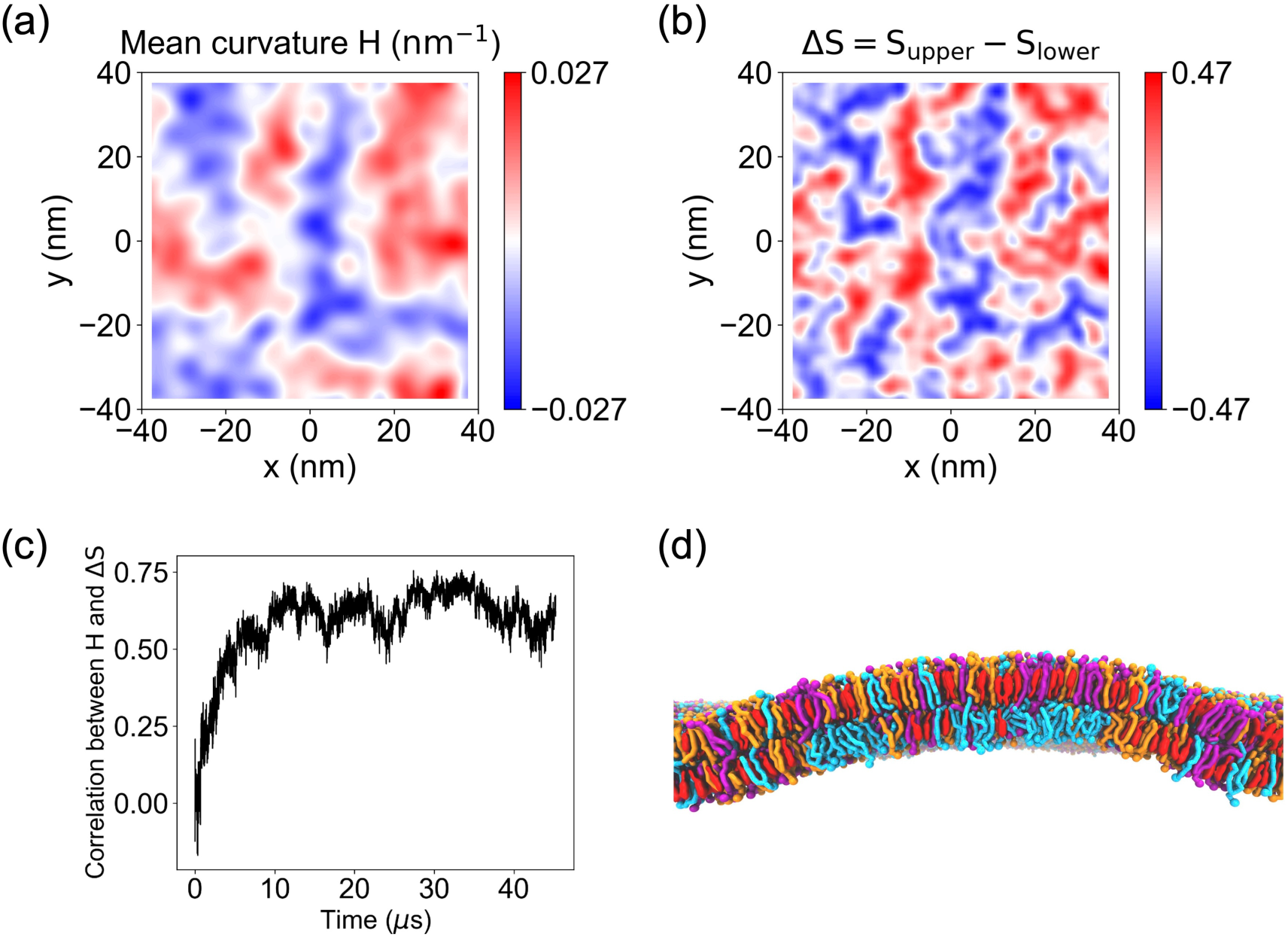
Coupling between anti-registered domains and curvature in the scrambled PM. (a) Two-dimensional distribution of the mean curvature in the scrambled PM, averaged over the 10 ns trajectory segment containing the frame with the largest root-mean-square height fluctuation of the bilayer midplane. (b) Two-dimensional distribution of the order parameter difference between the upper and lower leaflets, Δ*S* = *S*_upper_− *S*_lower_, in the scrambled PM, averaged over the same 10 ns segment as in panel a. (c) Time evolution of the Pearson correlation coefficient between mean curvature and Δ*S* in the scrambled PM. (d) Representative side-view snapshot of a positively curved region in the scrambled PM. Color codes are as follows: sphingomyelins, purple; mono-or di-unsaturated phosphatidylcholines, orange; polyunsaturated lipids, cyan; cholesterol, red.

Figure 5d shows a representative side view of a positively curved region in the scrambled PM. In this region, a highly ordered domain enriched in cholesterol and SMs in the upper leaflet faces a disordered domain enriched in polyunsaturated lipids in the lower leaflet. A previous experimental study suggested that ordered domains composed of SM and cholesterol have a slightly negative spontaneous curvature.^34^ The observed coupling implies that the disordered domains have an even more negative preferred curvature, likely reflecting the conical shape of polyunsaturated lipids. The resulting mismatch in preferred curvature between the two leaflets thus induces or amplifies membrane undulations.

The observed coupling between lipid distribution and membrane curvature is consistent with the mechanism termed “diffusional softening” in a previous study,^35–37^ whereby curvature-composition coupling lowers the apparent bending rigidity of mixed lipid membranes. That study, however, was limited to simple two-or three-component lipid mixtures. Our results suggest that diffusional softening also operates in a compositionally complex PM model, where it is mediated by anti-registered domains, underscoring the relevance of this mechanism to biological membranes. Why, then, were such large-scale curved structures absent in the asymmetric PM? Under the periodic boundary conditions, the membrane must maintain zero net mean curvature over its entire surface, so that positively curved regions must be balanced by negatively curved regions. The asymmetric PM is poorly suited to such deformation, mainly because its outer leaflet is almost uniformly ordered and nearly devoid of polyunsaturated lipids, which have a strongly negative spontaneous curvature; consequently, the transbilayer mismatch in preferred curvature is nearly uniform across the membrane plane and cannot be relieved by laterally alternating regions of opposite curvature. This constraint is not merely an artifact of the periodic boundary conditions: the PM of a living cell is large enough to be regarded as macroscopically flat on the length scales considered here, and the underlying cortical cytoskeleton may further help maintain this planar geometry.

### Long-Chain Sphingomyelins are Required for Domain Anti-Registration

Next, to identify the lipid species associated with domain anti-registration and local membrane curvature, we calculated the Pearson correlation coefficients between the density of each lipid species and both Δ*S* and the mean curvature of the bilayer midplane (Figure 6a). Cholesterol and SMs, the major components of the highly ordered domains, were positively correlated with both Δ*S* and mean curvature, indicating that they are enriched in positively curved regions. In contrast, polyunsaturated lipids, the primary components of the disordered domains, were negatively correlated with both quantities and are thus enriched in negatively curved regions. Mono- and di-unsaturated phosphatidylcholines exhibited near-zero correlations, indicating no clear curvature preference.

**Figure 6.**
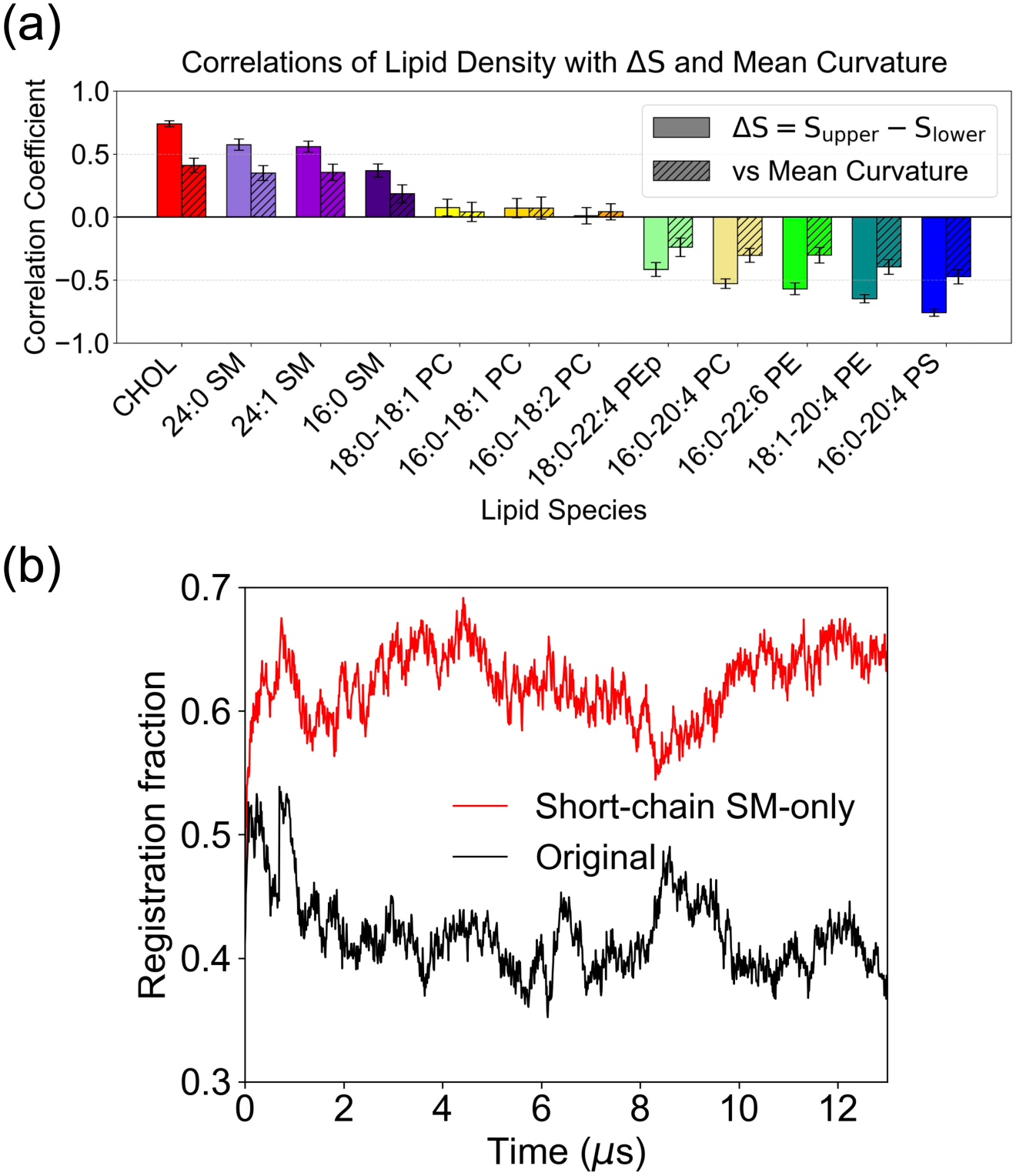
Long-chain sphingomyelins promote anti-registered domains. (a) Pearson correlation coefficients of lipid density with the interleaflet order parameter difference Δ*S* and mean curvature for each lipid species in the scrambled PM. Solid and hatched bars represent correlations with Δ*S* and mean curvature, respectively. Values were averaged over the final 10 μs. (b) Registration area fraction calculated from a 13 μs simulation of the scrambled PM in which all long-chain SMs (24:0 SM and 24:1 SM) were replaced with 16:0 SM (red curve). The black curve shows the registration area fraction of the original scrambled PM (Figure 4) for the first 13 μs.

Importantly, the long-chain SMs (24:0 SM and 24:1 SM) showed stronger positive correlations than 16:0 SM, indicating that long-chain SMs are preferentially located in anti-registered domains. Our previous CG MD study of phase-separated model membranes showed that long-chain SMs promote domain anti-registration by penetrating into the opposing leaflet and excluding cholesterol from the corresponding region.^38^ To examine whether such interdigitation also occurs in the scrambled PM model, we analyzed the probability density distributions along the *z*-axis of the terminal CG particles of the SM acyl chains and of cholesterol in a small scrambled PM model with a side length of 8.5 nm, in which no phase separation occurred (Figure S1). The acyl chains of 24:0 SM and 24:1 SM extended deeply into the opposing leaflet, partially overlapping with the cholesterol ring structure along the *z*-axis.

To confirm that long-chain SMs are also the major determinant of domain anti-registration in the scrambled PM, we performed an additional 13 μs simulation of the 76 nm scrambled PM in which all long-chain SMs were replaced with 16:0 SM, so that 16:0 SM was the only SM species. In this system, the registration area fraction (Figure 6b), calculated from the order parameter distributions (Figure S2), was approximately 0.62, indicating that the domains preferentially adopted a registered configuration. Thus, long-chain SMs are required for domain anti-registration in the scrambled PM.

Taken together, these results so far show that membrane curvature couples to anti-registered domains, in which ordered domains rich in long-chain SMs and cholesterol face disordered domains rich in polyunsaturated lipids, while domain enlargement allows the resulting undulations to develop over large length scales. The resulting convex and concave regions may lead directly to budding and invagination or may promote membrane remodeling indirectly by serving as platforms for curvature-sensing proteins. Previous studies have shown that polyunsaturated lipids facilitate membrane deformation,^39,40^ consistent with our observations. Our results further suggest that long-chain SMs may also be key determinants of membrane deformation. Because long-chain SMs are not rare lipids but are abundant in various cell types,^41^ this role may be relevant to membrane remodeling in many cellular contexts.

### System-Size Dependence of Domain Registration and Anti-Registration

We also found that the interleaflet domain organization of the scrambled PM depends strongly on system size. To examine whether the behavior observed in the large system (side length of approximately 76 nm) is reproduced in a smaller system, we simulated a scrambled PM with a side length of 34 nm. Unexpectedly, the domain organization across the two leaflets changed qualitatively with system size.

Figure 7a shows the two-dimensional order parameter distributions of the 34 nm scrambled PM with the original lipid composition after 20 μs of simulation. In contrast to the 76 nm system, domains of the same phase faced each other across the two leaflets, and the registration area fraction (black curve in Figure 7c) shows that this system remained registered throughout the simulation. The factors that destabilize anti-registered domains in the smaller system are discussed below.

**Figure 7.**
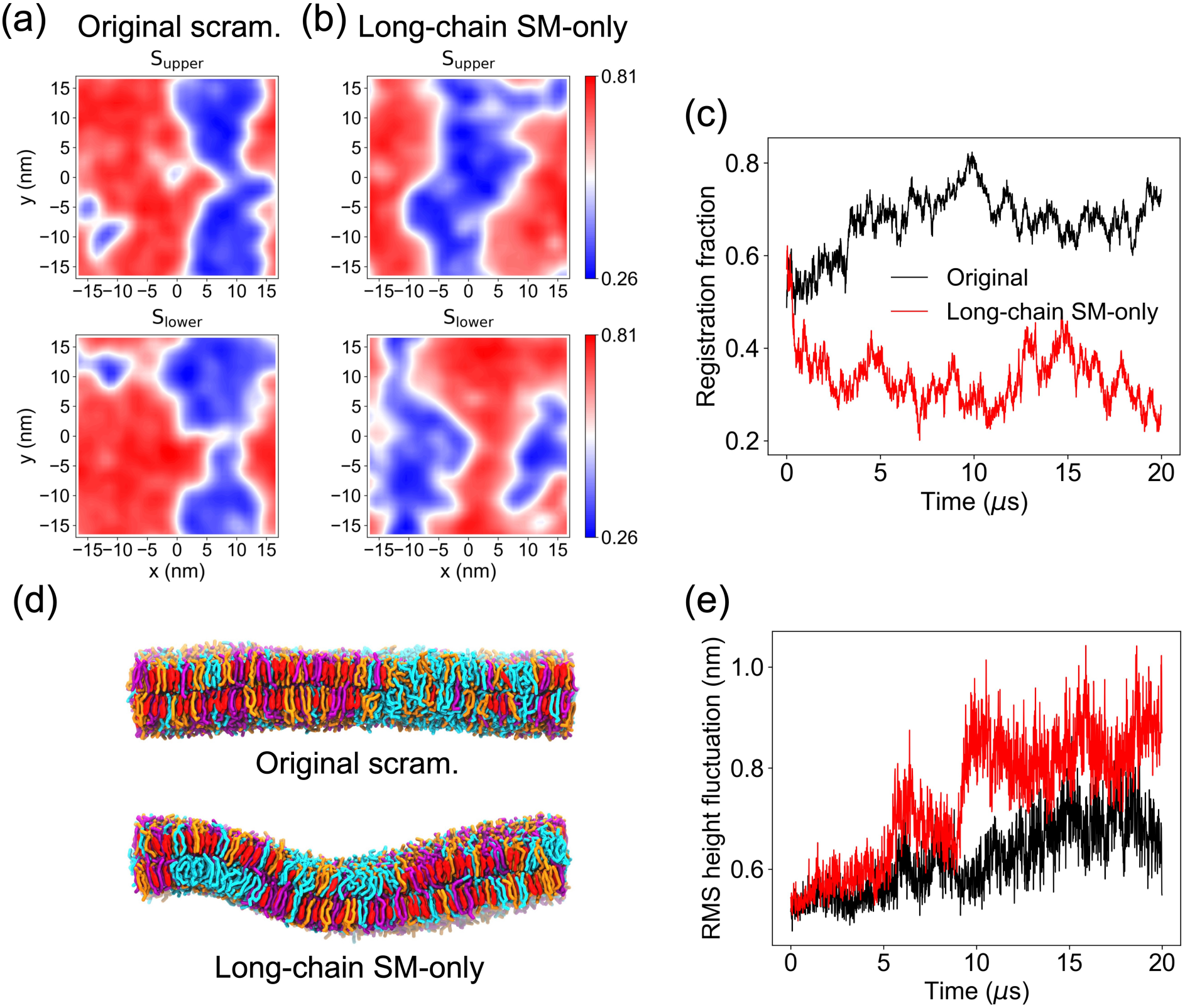
Domain organization and membrane undulations in small, scrambled PM systems with a side length of 34 nm. (a) Order parameter maps of the scrambled PM with the original lipid composition, calculated from the final 10 ns of the trajectory. (b) Order parameter maps of the scrambled PM in which the SM fraction consisted of equimolar 24:0 SM and 24:1 SM, with 16:0 SM omitted (long-chain SM-only system), calculated in the same manner. (c) Time evolution of the registration area fraction in the two systems. (d) Side-view snapshots of the final configurations of the two systems. Color codes are as follows: sphingomyelins, purple; mono- or di-unsaturated phosphatidylcholines, orange; polyunsaturated lipids, cyan; cholesterol, red. (e) Time evolution of the RMS height fluctuations of the two systems.

We first examined whether a high concentration of long-chain SMs can induce anti-registered domains even in the small box. For this purpose, we simulated a 34 nm system in which the SM fraction consisted of equimolar 24:0 SM and 24:1 SM, with 16:0 SM omitted. The order parameter distributions (Figure 7b) and registration area fraction (red curve in Figure 7c) show that ordered and disordered domains were anti-registered in this system. The side-view snapshots (Figure 7d) and RMS height fluctuations (Figure 7e) show that the system with the original lipid composition remained relatively flat, whereas the long-chain-SM-only system was strongly bent. These results highlight the strong ability of long-chain SMs to induce anti-registered domains and membrane curvature. It should be noted, however, that replacing 16:0 SM with long-chain SMs also increases the thickness of the ordered domains and hence the hydrophobic thickness mismatch between ordered and disordered domains. Because an anti-registered arrangement reduces the variation in total bilayer thickness across the membrane plane, a larger thickness mismatch could by itself favor anti-registration. The present comparison therefore cannot fully separate the contribution of the interleaflet interdigitation of long-chain SMs (Figure S1) from that of the enhanced hydrophobic mismatch and both effects may act together to stabilize anti-registered domains.

Why does the scrambled PM with the original composition adopt a registered rather than an anti-registered configuration when the system size is reduced? The relative stability of the two configurations is determined by several contributions, including the line tension at domain boundaries, the interfacial tension at the bilayer midplane, and the interdigitation of long-chain SMs described above. Because the perimeter-to-area ratio of a domain increases as the domain shrinks, a reduction in domain size would shift the balance among these contributions. However, the order parameter distributions (Figures 7a and 7b) and their spatial correlation functions (Figure S3) show that percolated domains comparable in size to those in the 76 nm system form even in the 34 nm system, although they percolate across the periodic boundaries. Because the shape and boundary length of such percolated domains may be affected by the box geometry, a contribution from line tension cannot be completely ruled out. Nevertheless, the comparable domain sizes suggest that these contributions alone are unlikely to account for the observed size dependence, which led us to also consider a factor that depends on the box size itself rather than on the domain size.

In the small system, the finite box restricts the wavelength of undulations. In the large system, by contrast, this degree of freedom is far less constrained, and we assume that the membrane can therefore adopt a curvature close to the spontaneous curvature of anti-registered domains. Because long-chain SMs are enriched in the anti-registered regions of the original large system, anti-registered domains in the long-chain-SM-only small system are expected to share a similar spontaneous curvature.

To compare the magnitudes of the mean curvature in these two systems, we plotted the mean curvature against Δ*S* and fitted the data with a linear function (Figure S4). The long-chain SM-only small system spanned a wider range of curvature and showed a steeper slope than the large system with the original composition, indicating stronger bending. This suggests that, because the wavelength of undulations is restricted, the small membrane much bend beyond the spontaneous curvature of the anti-registered domains, thereby incurring an elastic penalty. In the long-chain-SM-only small system, the free-energy gain from anti-registration outweighs this penalty; with the original composition, however, the smaller gain associated with the lower long-chain SM content cannot compensate for it, and the small system therefore remains registered. Alternatively, if the spontaneous curvature of the anti-registered domains increases with their long-chain SM content, the spontaneous curvature for the original composition may simply be too small to be compatible with undulations whose wavelengths fit within the 34 nm box.

## CONCLUSION

Our CG-MD simulations of asymmetric and scrambled PM models showed that the asymmetric PM remained nearly flat, whereas the scrambled PM exhibited substantially larger membrane undulations. These enhanced undulations were associated with lipid scrambling-induced domain reorganization characterized by two key features: domain anti-registration and domain enlargement. In the anti-registered configuration, ordered domains rich in cholesterol and SMs face disordered domains rich in polyunsaturated lipids across the bilayer. As the disordered domain have a more negative preferred curvature, this arrangement couples domain organization to membrane curvature. Domain enlargement, in turn, allows these undulations to extend over large length scales. Anti-registration is stabilized by long-chain SMs, whereas domain enlargement is likely driven by the coexistence of high concentrations of SMs and polyunsaturated lipids within the same leaflet, a condition absent in the asymmetric state. Simulations of smaller systems further revealed that domain registration depends strongly on system size, a finite-size effect that should be taken into account in simulations of compositionally complex membranes. These simulations also confirmed the strong ability of long-chain SMs to induce anti-registered domains and membrane curvature. Although the present models represent the fully scrambled limit and lack membrane proteins and the cortical cytoskeleton, these findings provide a molecular-level physical basis for understanding how lipid scrambling promotes large-scale membrane remodeling and suggest a previously unrecognized role for long-chain SMs in coupling interleaflet interactions to membrane curvature.

## SUPPORTING INFORMATION

Lipid compositions of all simulated scrambled PM models (Table S1); probability density distributions along the bilayer normal of the terminal CG particles of the SM acyl chains and of cholesterol in a small scrambled PM, and CG mapping of cholesterol in the SPICA FF (Figure S1); two-dimensional order parameter distributions of the 76 nm scrambled PM in which all long-chain SMs were replaced with 16:0 SM (Figure S2); spatial correlation functions of the order parameter in the 34 nm scrambled PMs (Figure S3); mean curvature as a function of the interleaflet order parameter difference in the 76 nm scrambled PM with the original composition and in the 34 nm long-chain-SM-only scrambled PM (Figure S4) (PDF)

## ACKNOWLEDGMENTS

This work was supported by JST SPRING, Japan Grant Number JPMJSP2126, and JSPS KAKENHI Grant Numbers JP24H00843, JP25K01729, and JP26K22773. Calculations were performed on the supercomputer facilities of the Institute for Solid State Physics, the University of Tokyo (ISSPkyodo-SC-2025-Ea-0002, 2025-Eb-0001, and 2026-Ea-0003), and the Research Center for Computational Science, Okazaki, Japan (Project: 25-IMS-C094 and 26-IMS-C087).

## Table of Contents Image

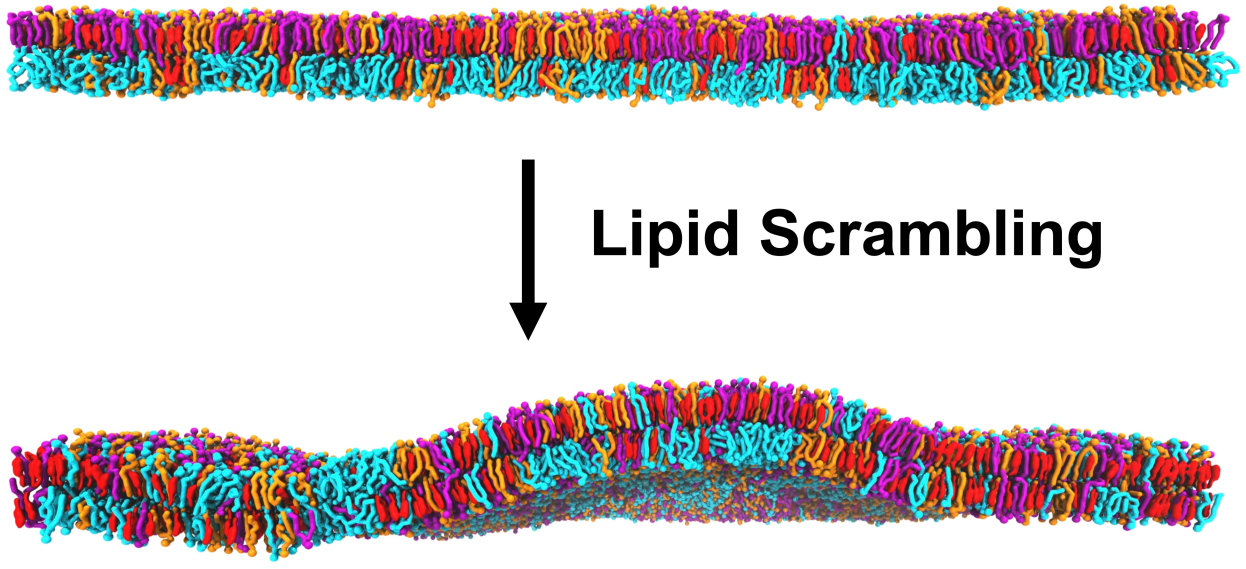

