## Supporting Information for "Lipid Scrambling Amplifies Plasma Membrane Undulations via Anti-Registered Domains Stabilized by Long-Chain Sphingomyelin"

**Table S1.** Lipid compositions of the simulated scrambled PM models. The number of molecules of each lipid species, summed over the upper and lower leaflets, is listed for each system.

| **Lipid** | Original (76 nm) | Short-chain -SM-only (76 nm) | Original (34 nm) | Long-chain -SM-only (34 nm) | Original (8.5 nm) |
|---|---|---|---|---|---|
| Cholesterol | 10530 | 10530 | 2080 | 2080 | 130 |
| 16:0 SM | 1458 | 3564 | 288 | 0 | 18 |
| 24:1 SM | 1134 | 0 | 224 | 352 | 14 |
| 24:0 SM | 972 | 0 | 192 | 352 | 12 |
| 18:0-18:1 PC | 810 | 810 | 160 | 160 | 10 |
| 16:0-18:1 PC | 648 | 648 | 128 | 128 | 8 |
| 16:0-18:2 PC | 3240 | 3240 | 640 | 640 | 40 |
| 16:0-20:4 PC | 486 | 486 | 96 | 96 | 6 |
| 16:0-20:4 PS | 2430 | 2430 | 480 | 480 | 30 |
| 16:0-22:6 PE | 1458 | 1458 | 288 | 288 | 18 |
| 18:1-20:4 PE | 648 | 648 | 128 | 128 | 8 |
| 18:0-22:4 PEp | 1620 | 1620 | 320 | 320 | 20 |
| **Total** | **25434** | **25434** | **5024** | **5024** | **314** |

Lipid abbreviations are as follows: SM, sphingomyelin; PC, phosphatidylcholine; PS, phosphatidylserine; PE, phosphatidyl ethanolamine; PEp, PE plasmalogen.

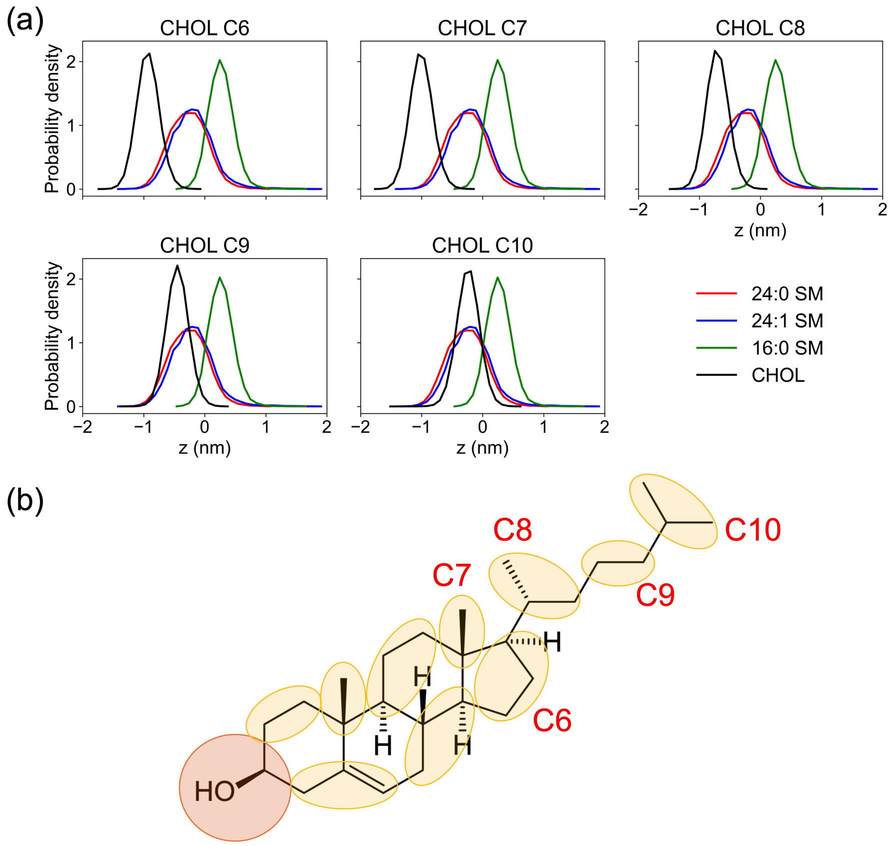


**Figure S1.** (a) Probability density distributions along the *z*-axis of the terminal CG particles of the SM acyl chains and of cholesterol in a small scrambled PM without phase separation (side length of 8.5 nm). The distributions were calculated from the final 200 ns of a 300 ns simulation. (b) CG mapping of cholesterol in the SPICA FF. The CG particle names correspond to the titles of the individual panels in panel a.

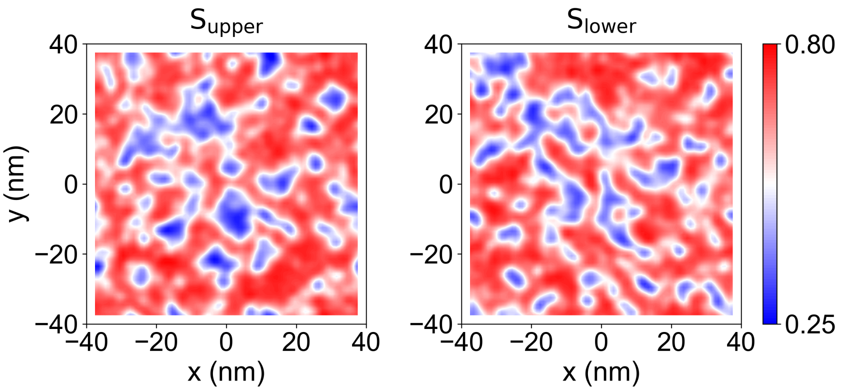


**Figure S2.** Two-dimensional order parameter distributions of the 76 nm short-chain-SM-only scrambled PM, in which all long-chain SMs were replaced with 16:0 SM, at the end of the 13 μs simulation (see Figure 6b).

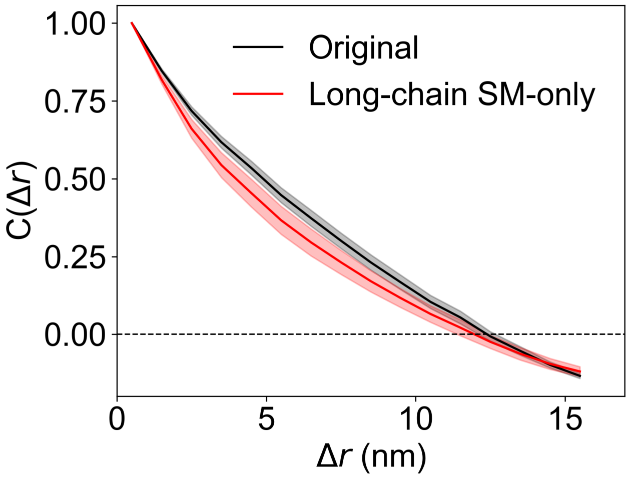


**Figure S3.** Spatial correlation functions of the order parameter for the 34 nm scrambled PM models with the original lipid composition and with the long-chain-SM-only composition.

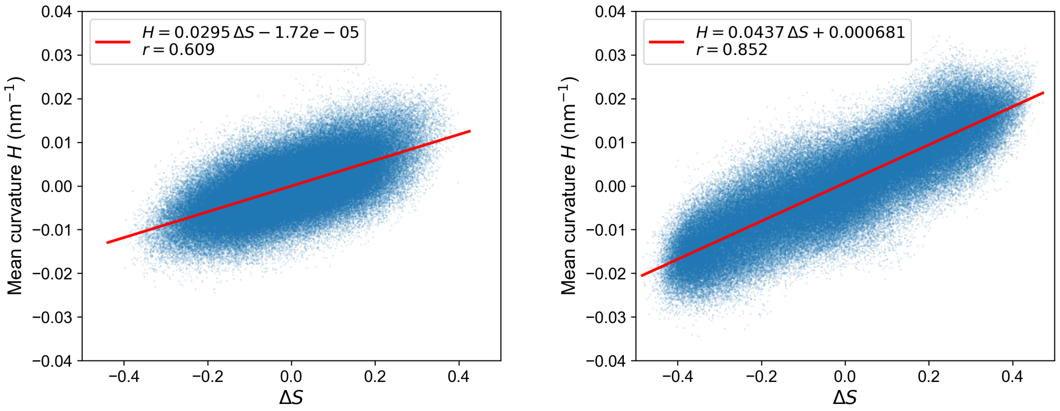


**Figure S4.** Mean curvature as a function of the interleaflet order parameter difference, $\Delta S$, for the 76 nm scrambled PM with the original lipid composition (left) and the 34 nm long-chain-SM-only scrambled PM (right). Red lines indicate linear fits.
